# Tardigrades dramatically upregulate DNA repair pathway genes in response to ionizing radiation

**DOI:** 10.1101/2023.09.07.556677

**Authors:** Courtney M. Clark-Hachtel, Jonathan D. Hibshman, Tristan De Buysscher, Bob Goldstein

**Author notes:** Biology Department, The University of North Carolina at Asheville, Asheville, NC 28804, USA.

## Abstract

Tardigrades can survive remarkable doses of ionizing radiation, up to about 1000 times the lethal dose for humans. How they do so is incompletely understood. We found that the tardigrade *Hypsibius exemplaris* suffers DNA damage upon gamma irradiation, but damage is repaired. We show that tardigrades have a specific and robust response to ionizing radiation: irradiation induces a rapid, dramatic upregulation of many DNA repair genes. By expressing tardigrade genes in bacteria, we validate that increased expression of some repair genes can suffice to increase radiation tolerance. We show that at least one such gene is necessary for tardigrade radiation tolerance. Tardigrades’ ability to sense ionizing radiation and massively upregulate specific DNA repair pathway genes may represent an evolved solution for maintaining DNA integrity.

**One-Sentence Summary:** Tardigrades exposed to ionizing radiation survive DNA damage by massively upregulating DNA repair pathway genes.

## Main Text

Some organisms have evolved to survive conditions that to most organisms would be lethal, including extreme heat, extreme cold, and desiccation (*1–7*). Revealing the mechanisms that these organisms employ to survive under stressful conditions can aid in understanding stress tolerance and may contribute to improving the survival of less tolerant organisms, cells, or biological materials in the face of stress.

Tardigrades are well known for their ability to survive in environments where other animals would not (*5*, *6*). Some tardigrade species have been demonstrated to survive desiccation as well as extreme temperatures, pressures, and levels of ionizing radiation (IR) (*5–11*). For example, while the dose of IR at which 50% of humans would die (LD50) is 5 gray (Gy), the tardigrade *Hypsibius exemplaris* can survive ~4,000 Gy (*12*). At these levels of IR we would expect massive amounts of DNA damage and genomic instability (*13*, *14*).

Little is known about the specific mechanisms that underly tardigrade extreme resistance to genotoxic stress. Most of what is known comes from work in the tardigrade *Ramazzottius varieornatus*, a species with a similar IR tolerance to *H. exemplaris* (*14*). *R. varieornatus* produces a DNA damage suppressing protein (Dsup) that can confer IR resistance when expressed in human cells (*14*, *15*). Biochemical studies of this protein have revealed that it protects DNA from IR by binding to DNA and nucleosomes and protecting DNA from hydroxyl radicals that are generated by IR-exposed cells (*16*). The identification of Dsup suggested that tardigrades can survive high doses of IR through the employment of protective mechanisms that prevent damage to the DNA. However, it remains unclear if protective mechanisms can fully explain the extreme IR tolerance of tardigrades. The protein sequence of Dsup is not well conserved in tardigrades; hence, it is unclear if other tardigrade species’ Dsup proteins have the same protective abilities (*15*).

Here, we set out to understand how the tardigrade *H. exemplaris* can survive extreme IR. Through DNA damage assays, expression analyses, and functional studies, we show that *H. exemplaris* tardigrades do experience DNA damage upon IR exposure, that they upregulate DNA repair transcripts to a remarkable and unexpected degree in response to IR, and that the increased expression of some DNA repair transcripts is both sufficient to confer IR tolerance to bacteria and necessary for tardigrade IR tolerance.

## Results

### H. exemplaris experiences DNA damage from ionizing radiation

To visualize the level and location of DNA damage in tardigrades following IR exposure, we adapted a terminal deoxynucleotidyl transferase dUTP nick end labeling (TUNEL) assay for use on whole animals (see Materials and Methods). TUNEL assays are commonly used to visualize DNA single-stranded (ss) and double-stranded (ds) breaks (*17*). We exposed animals to a well-tolerated dose of gamma irradiation (2,180 Gy), as well as a dose near the LD50 (4,360 Gy) (*12*). Animals that were never exposed to IR had little TUNEL signal in their nuclei (Fig. 1). We found that animals exposed to IR had significantly more TUNEL signal per nucleus than control animals (Fig. 1). This suggests that the tardigrades indeed experience DNA damage from IR exposure. To determine if damage is repaired, we exposed animals to the same doses of gamma irradiation and allowed them to recover for 24 hours (Fig. 1). After a sub-lethal irradiation dose, animals that were exposed to IR and allowed to recover showed a significant reduction in TUNEL signal per nucleus over 24 hours of recovery (Fig. 1D). These results suggest that *H. exemplaris* experiences DNA damage upon extreme levels of IR but is then able to repair much of the damage.

**Fig. 1.**
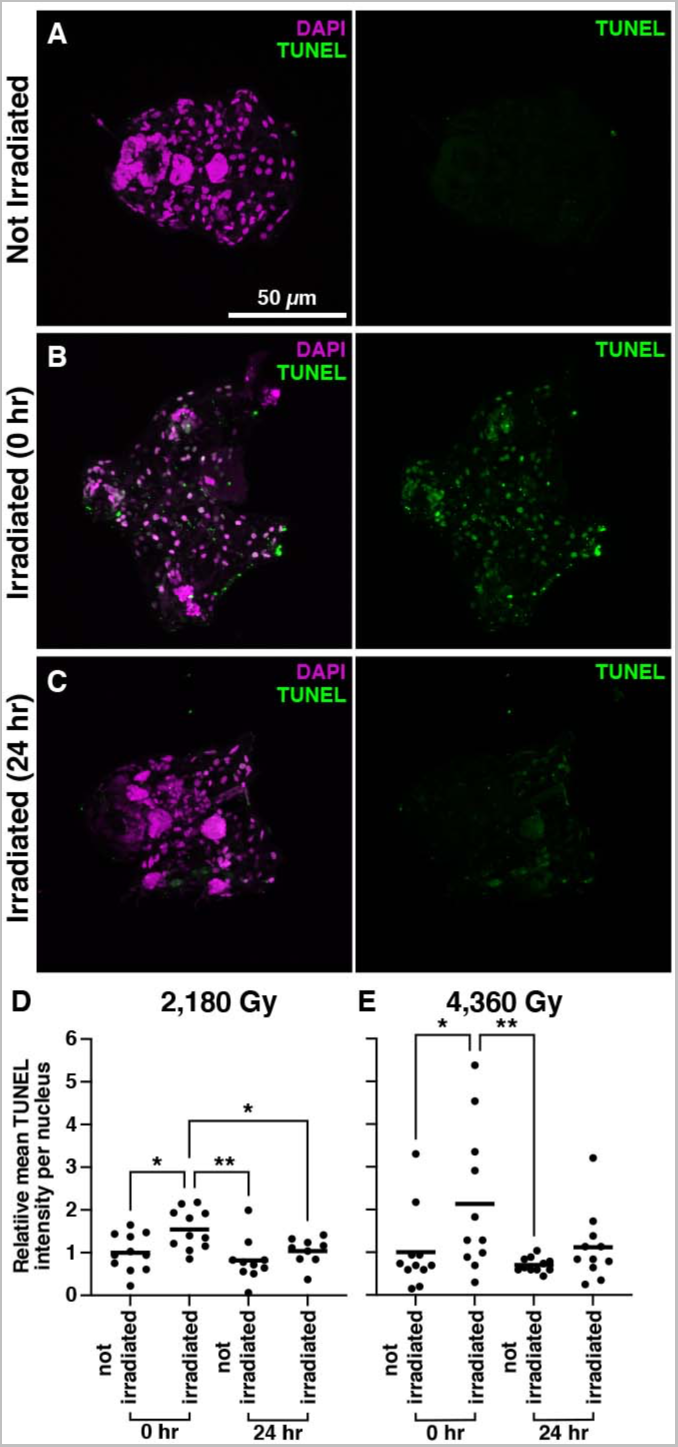
Visualizing DNA damage in tardigrades following ionizing radiation exposure. (**A-C**) Representative images of TUNEL signal in individuals that were not exposed to IR (**A**), exposed to 4,360 Gy IR (**B**), and exposed to 4,360 Gy IR and allowed to recover for 24 hours (**C**). Scale bar applies to all images. Animals were physically disrupted for TUNEL protocol so above images are fragments of whole adults. Anterior is to the left. (**D and E**) Plots displaying the relative mean intensity of TUNEL signal per nucleus for 2,180 Gy (**D**) and 4,360 Gy (**E**) IR exposure. For each plot the groups from left to right are as follows: not exposed to IR, exposed to IR, not exposed to IR and left for 24 hours, exposed to IR and allowed to recover for 24 hours, (n=11 individuals for all groups, except 2,180 Gy not irradiated 24 hr (n=10) and 2,180 Gy irradiated 24 hr (n=9)). A one-way ANOVA followed by a post-hoc Dunnett test to the mean of the irradiated timepoint 0 for each experiment was used to determine significant differences between treatment groups. Significance is as follows: *p<.05 ** p<.01. 2,180 Gy 0 hr vs. not irradiated 0 hr: p=.02, vs. not irradiated 24 hr: p=.002, and vs irradiated 24 hr: p=.04. 4,360 Gy 0 hr vs vs not irradiated 0 hr: p=.04 and vs not irradiated 24 hr: p=.008.

### Tardigrades massively upregulate the transcription of DNA repair pathway genes after exposure to ionizing radiation

*H. exemplaris* could be engaging in a variety of measures to compensate for DNA damage, including transcriptomic responses. The well-established DNA damage response only minimally involves transcriptional responses (*18*), yet some modest transcriptional responses to DNA damaging agents have been observed in other animals with a typical enrichment of 1.5-4 fold for any responsive transcript (*19–25*). To examine tardigrade transcriptomes after IR, we performed messenger RNA sequencing (mRNA-Seq) on animals after exposure to 100, 500, or 2,180 Gy doses of IR. *H. exemplaris* can survive and reproduce after exposure to 100 Gy. After exposure to 500 or 2,000 Gy (about half of the LD50), they survive well but no longer reproduce (*12*).

Differential expression analysis revealed that tardigrades have a robust transcriptional response to IR exposure, with 4,590 transcripts significantly upregulated and 4,687 downregulated in response to 500 Gy IR (p<.05, Fig. 2A). We were intrigued to find that 7 of the top 15 most significantly enriched transcripts encoded proteins of DNA repair pathways (Fig. 2A, Table S1). These transcripts included representatives from Base Excision Repair (BER) (*DNA LIG1*, *PNKP*, *PARP3*, *PARP2*, and *PCNA*) and Non-Homologous End Joining (NHEJ) (*XRCC5*, which encodes Ku80, and *DNA LIG4*) (Fig. 2A, Table S1), all of which were upregulated more than 32-fold (Table S1). By comparison, a recent study of the transcriptional response to IR in mammalian cells identified only *PCNA* and *LIG1* from this list, both of which were upregulated less than 2-fold (*20*). The remaining genes from the top 15 list are predicted to encode two tardigrade-specific proteins with no conserved domains, two predicted histone proteins, a mitochondrial chaperone BCS1, a protein phosphatase 1B, a protein with RING-HC and WWE domains, and a partial Ku70 protein with no predicted DNA repair function (see Materials and Methods (*26*)) (Table S1). The fact that multiple DNA repair pathway transcripts are represented in the most significantly enriched transcripts indicates that *H. exemplaris* responds to the damage caused by IR by upregulating genes encoding proteins that can correct damage. The degree of upregulation after IR was high (32- to 315-fold). In addition, DNA repair genes constituted some of the most highly represented transcripts in the animals’ transcriptome after IR (Fig 1B-D and Table S1 and S2) bringing some DNA repair transcripts up nearly to the level of highly expressed housekeeping genes like elongation factor 1-alpha and cytoplasmic actin (determined by TPM, Table S2). The pathways that were most represented here (NHEJ and BER) are most apt to repair the types of DNA damage that commonly result from exposure to IR (*27*). IR can directly generate dsDNA breaks, which are repaired primarily by the NHEJ pathway (*27*, *28*). IR exposure can also lead to the production of reactive oxygen species, which can cause ssDNA breaks as well as damaged bases, both of which are repaired by BER (*27*, *29*).

**Fig. 2.**
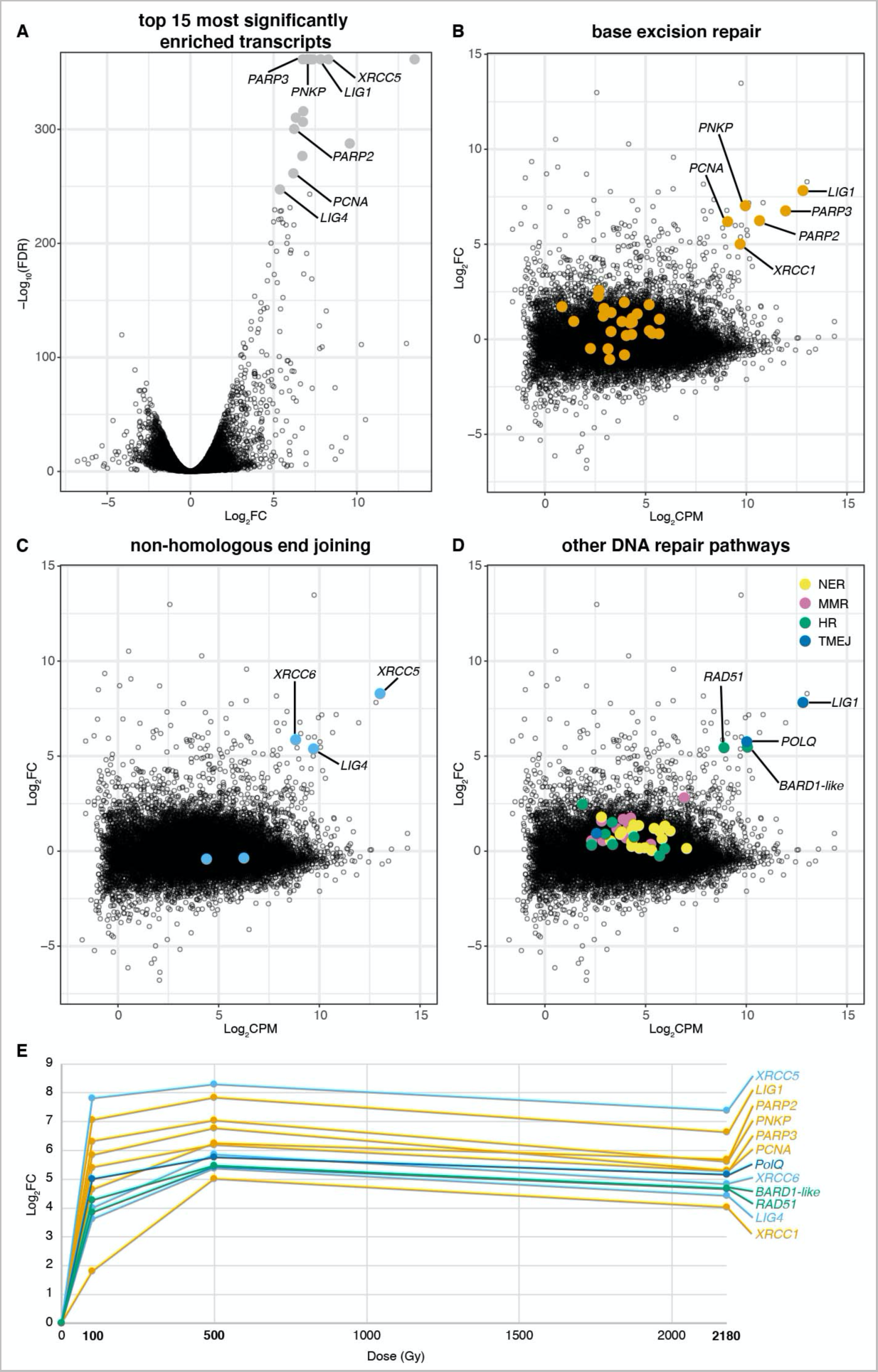
Tardigrades dramatically increase expression of certain DNA repair transcripts in response to ionizing radiation. (**A**) Volcano plot of Log_2_FC by −Log_10_(FDR) showing the transcriptional response of *H. exemplaris* to 500 Gy IR. The top 15 most significantly enriched transcripts (by FDR) are marked in gray. DNA repair pathway genes among the top 15 are labeled. (**B-D**) MA plots displaying Log_2_FC of *H. exemplaris* transcripts in response to 500 Gy IR with (**B**) transcripts encoding BER proteins marked in orange, (**C**) transcripts encoding NHEJ proteins marked in light blue, and (**D**) transcripts encoding NER, MMR, HR, and TMEJ proteins marked in yellow, pink, green, and dark blue, respectively. Transcripts encoding DNA repair proteins that are significantly enriched are indicated by name. Note that LIG1 functions in two pathways. (**E**) Plot showing Log_2_FC for enriched DNA repair transcripts at 100, 500, and 2180 Gy doses of IR. Colors are the same as in MA plots. LIG1 is colored as BER (orange), but also functions in TMEJ (dark blue)

We were curious if other transcripts from NHEJ and BER pathways or from other DNA repair pathways were also enriched after exposure to IR. We found that multiple BER pathway genes were indeed enriched following IR exposure (Fig. 2B, Table S1 and S3). In addition to the BER genes listed above, *XRCC1* was also enriched. We conclude that many of the genes important for BER are upregulated in response to IR. From NHEJ, *XRCC6* (which encodes Ku70) was also enriched following IR (Fig. 2C, Table S3) which, in combination with XRCC5 and *LIG4* mentioned above, forms a complete set of the minimal proteins sufficient to perform NHEJ repair *in vitro* (*30*).

To examine whether *H. exemplaris* upregulates other DNA repair pathways in response to IR, we also looked at transcript enrichment for genes from the Mismatch Repair (MMR, repairs base mismatches), Nucleotide Excision Repair (NER, removes bulky adducts), Homologous Recombination (HR, repairs dsDNA breaks), and Theta-Mediated End Joining (TMEJ, repairs dsDNA breaks) pathways (*27*, *31*, *32*). Amongst HR-associated genes, *RAD51* and *BARD1-like* were enriched following IR (Fig. 2D, Table S3). Transcripts encoding two out of the three homologs for TMEJ proteins that we identified in tardigrades (DNA polymerase Theta (POLQ) and LIG1) were also significantly enriched following IR (Fig 2D, Table S1 and S3). No genes from NER or MMR pathways had transcripts significantly enriched following IR (Fig 2D, Table S4). Taken together, these results reveal specificity in the transcriptional response of tardigrades to IR, with animals increasing the expression of DNA repair genes from pathways that deal with the types of damage expected to result from IR. We found that many of these genes are dramatically upregulated even after a 100 Gy dose over one hour (Fig. 2E), suggesting a rapid and robust response.

### DNA repair transcripts are upregulated throughout the animal following ionizing radiation exposure with some tissue-specific enrichment

Our results above, demonstrating a strong and diverse response to IR, led us to wonder whether these responses occur throughout entire tardigrades or whether there are specific tissues that drive this response. To determine whether specific tissues respond to IR by upregulating repair transcripts, we performed *in situ* hybridization for a sample of the DNA repair transcripts that were enriched following IR exposure. After exposure to 100 Gy IR, enrichment of transcripts was detectable via *in situ* hybridization for the DNA repair transcripts that we examined, confirming our mRNA-Seq results (Fig. 3, Fig. S1–S3). All DNA repair transcripts that we observed became enriched in nearly all examined tissues after IR exposure (Fig. S3) but also demonstrated some extent of tissue-specific enrichment (Fig. 3, Fig. S1–S3). For multiple DNA repair genes, transcripts were especially enriched in secretory tissues (salivary glands and claw glands) and the hindgut (arrow, arrowhead, and dashed outline in Fig. 3, Fig. S1 and S2, respectively, Fig. S3 (*33*)). In addition, we observed expression enrichment in storage cells (coelomocytes) for all but one of the DNA repair transcripts we observed (Fig. S3). We conclude that the responses to IR exposure that we have identified are strongest in certain tissues, including secretory tissues, which are expected to be especially active in transcription and translation (*34*).

**Fig. 3.**
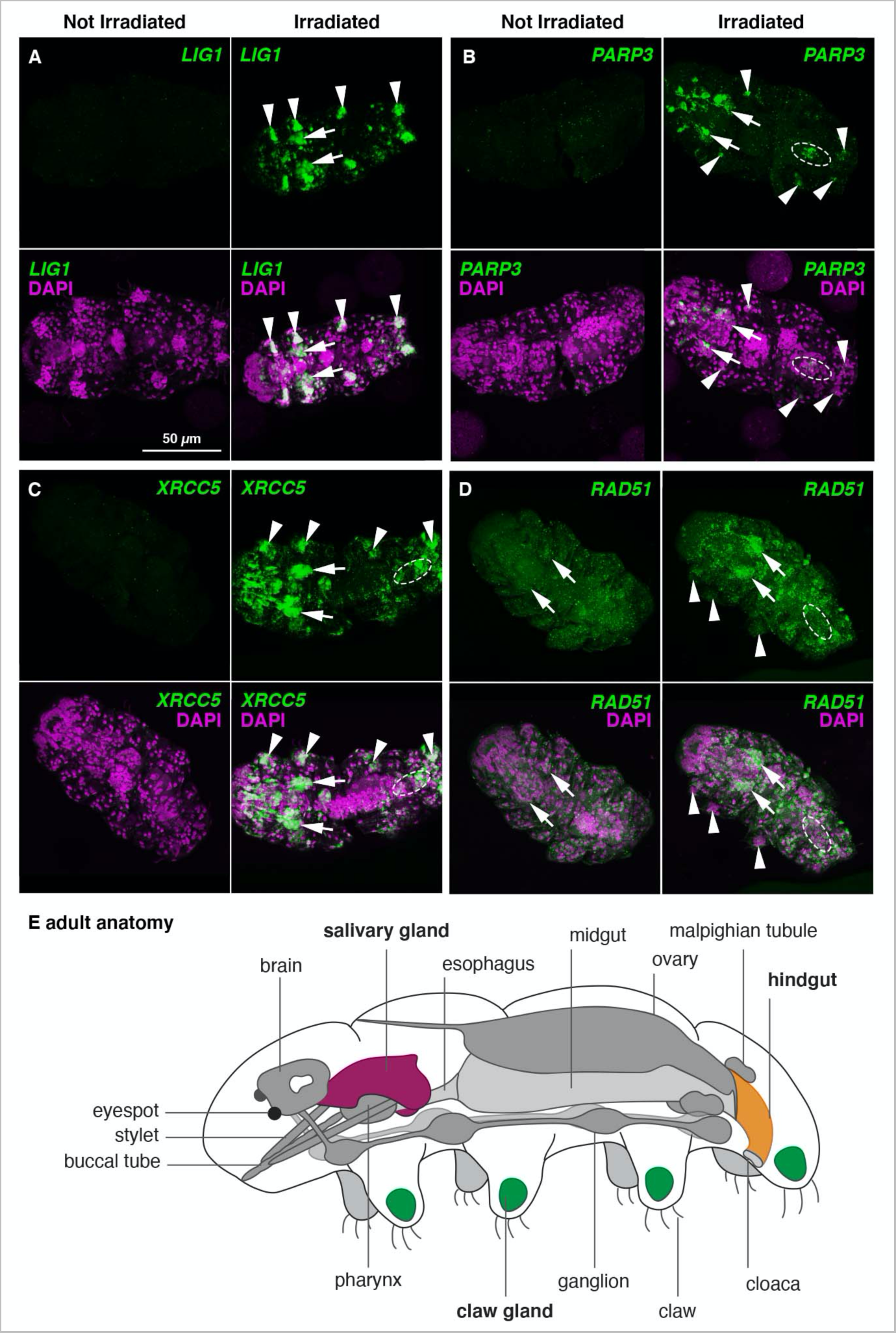
Tissue-specific enrichment of tardigrade DNA repair transcripts following ionizing radiation exposure. (**A-D**) Representative images of *in situ* hybridization for DNA repair transcripts with and without exposure to 100 Gy ionizing radiation. Exposure and contrast were adjusted to visualize regions of most intense signal. Expression in salivary glands (arrows), claw glands (arrowheads), and hindgut (dashed outlines) is indicated where seen. Transcripts encoding members of the (**A**) TMEJ, (**A-B**) BER, (**C**) NHEJ, and (**D**) HR pathways are represented. Scale bar in A applies to all images. Anterior is to the left. (**E**) Schematic of a lateral view of an adult tardigrade with salivary glands (burgundy), claw glands (green), hindgut (orange), and other landmark structures (gray) indicated (adapted from (*44*)).

### Expression of tardigrade DNA repair transcripts in bacteria can confer resistance to ionizing radiation

We considered it likely that the increased expression of DNA repair transcripts that we found in *H. exemplaris* might be sufficient to protect against IR exposure. To test whether increased expression of these transcripts can be sufficient to increase protection against IR, we expressed tardigrade DNA repair genes heterologously in *Escherichia coli* (*E. coli*). Bacteria induced to express tardigrade DNA repair genes were exposed to 2,180 Gy IR to see if they would survive better than *E. coli* containing control expression vectors that were either empty (no gene insertion) or contained a control sequence encoding GFP. In addition, we used a vector expressing the *R. varieornatus Dsup* gene as a positive control, as it has been previously shown to improve radiation survival of human cells (*14*). We found that expression of some tardigrade DNA repair genes could significantly improve the IR tolerance of *E. coli* relative to controls (Fig. 4A and Fig. S4). Transcripts that improved survival included *RAD51*, *XRCC1*, *FEN1*, *LIG1, PARP2,* and *POLB*. All of these genes except for *RAD51* (HR pathway) encode proteins in the BER pathway (Fig. 4A). For some DNA repair genes, expression conferred about as strong protection as did expression of the known DNA protectant Dsup (Fig. 4A). Although the mechanisms of protection are unknown in these bacterial expression experiments, and some genes may protect bacteria by different mechanisms than used in tardigrades, these data reveal that increased expression of these transcripts, such as we found in tardigrades, can indeed be sufficient to confer increased protection against IR damage.

**Fig. 4.**
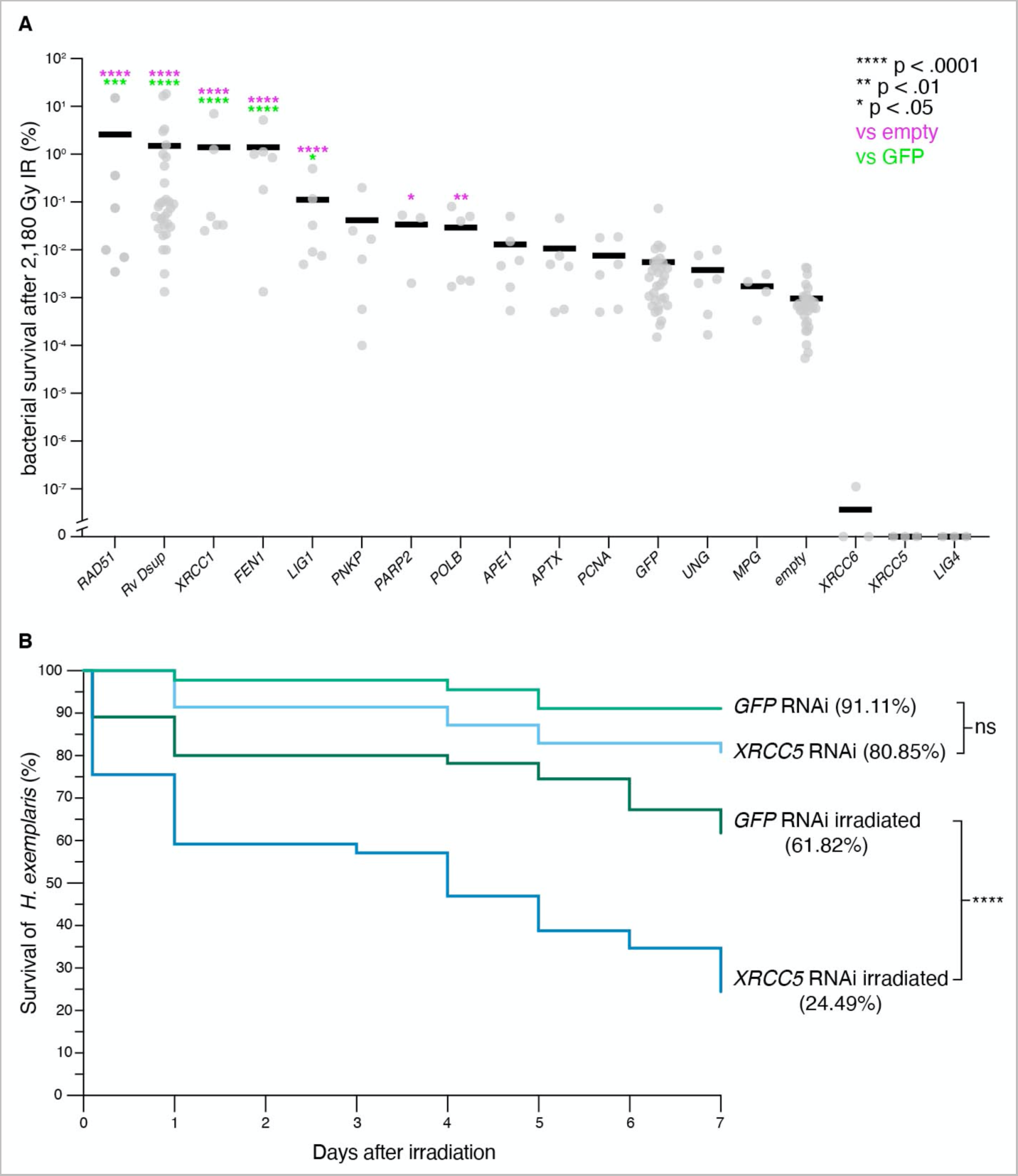
High levels of certain DNA repair genes are sufficient to improve bacterial survival after ionizing radiation exposures and necessary for tardigrade ionizing radiation survival. (**A**) Plot of tardigrade DNA repair transcripts organized from most efficient (left) to least efficient (right) at improving bacterial IR survival, including controls (GFP and empty). N=6 for all transcripts except for *Rv Dsup* (N=31), *PARP2* (N=3), *MPG* (N=4), GFP (N=31), empty (N=31), *XRCC6* (N=3), *XRCC5* (N=3), and *LIG4* (N=3). (**B**) Survival curves tracking the percent survival of animals following RNAi through 7 days after exposure to 2,180 Gy IR. Groups are as follows: *GFP* RNAi Control (light green, n=45), *XRCC5* RNAi Control (light blue, n=47), *GFP* RNAi irradiated (dark green, n=55), and *XRCC5* RNAi irradiated (dark blue, n=49). **** = p<.0001 (log-rank test).

### Increased expression of a DNA repair transcript is required for tardigrade ionizing radiation tolerance

To determine if high levels of a DNA repair transcript are essential for the ability of tardigrades to survive IR, we attempted to decrease the amount that a DNA repair transcript enriches in response to IR via RNA interference (RNAi). *H. exemplaris* is amenable to RNAi and has a systemic RNAi response: genes can be targeted in adults and their offspring by injection of double-stranded RNA (dsRNA) into individual animals (*35*, *36*). We chose *XRCC5* as a target because it was among the most significantly enriched gene transcripts following exposure to IR (Fig. 2A and C, Table S1 and S2). Most animals that were injected with dsRNA targeting either *XRCC5* or the control gene *GFP* survived over a 7-day period in the absence of IR exposure (Fig. 4B). After exposure to IR, animals that were injected with dsRNA targeting *XRCC5* experienced significantly more lethality than the *GFP*-targeted group (Fig. 4B). We conclude that the high levels of *XRCC5* transcripts that we found in tardigrades after exposure to IR contribute to the animals’ ability to survive this stress. This result suggests that at least one of the genes we identified as dramatically upregulated (enriched 315-fold) following IR plays a functional role in surviving IR exposure.

## Discussion

We found that the tardigrade *H. exemplaris* experiences DNA damage upon extreme doses of IR, and that they can repair much of that damage. Our mRNA-Seq analysis revealed an unexpectedly strong upregulation of DNA repair pathway genes in response to IR, with some transcripts enriched close to 300-fold, becoming among the most-represented transcripts in the animal’s transcriptome. The repair pathways that were most affected are those most clearly implicated in repairing the types of DNA damage that would be expected following IR exposure: BER, which repairs oxidative damage and ssDNA breaks, and NHEJ, which repairs dsDNA breaks. The specificity and magnitude of this transcriptional response suggests that tardigrades have mechanisms for sensing the DNA damage caused by IR and in response, dramatically increase the expression of specific DNA repair pathways. We found that RNAi targeting one such gene compromised the tardigrades’ ability to survive high doses of IR. This suggests that the transcriptional response we found through our mRNA-Seq analysis, at least for this example and possibly for the other genes, is important to tardigrade IR tolerance abilities. We also found that strong expression of some of these DNA repair transcripts alone is sufficient to confer IR tolerance to bacteria. We conclude that tardigrades have an adaptive response to DNA damage-inducing radiation that is unique to date: they survive the damage at least in part by massive transcriptional upregulation of DNA repair pathway genes. Taken together, these results expand our understanding of the mechanisms that animals use to maintain DNA integrity under damaging conditions and may provide potential new routes forward to improving DNA stability in other systems.

Why tardigrades have evolved strong IR tolerance is enigmatic, given that it is unlikely that tardigrades were exposed to high doses of IR in their evolutionary history. One possible explanation for their exceptional IR tolerance is that their adaptation for desiccation, a stress they likely experience frequently and can survive, has given them an ability to recognize and respond to DNA damage and hence a cross-tolerance to IR (*11*). Long-term desiccation can also result in genome instability and DNA damage (*37*, *38*). We investigated the transcriptome profile of tardigrades entering desiccation to see if DNA repair transcripts were also enriched (*36*, *39*). We did see a slight enrichment in some transcripts of the BER pathway (Fig. S5), but this enrichment was below log_2_FC of 2, not nearly as strong as the enrichment observed after IR exposure. While animals entering desiccation did not show strong enrichment of DNA repair transcripts, it remains possible that these transcripts could be robustly expressed later, upon rehydration. It remains enigmatic why tardigrades have evolved strong IR tolerance.

Transcriptional responses to IR have been interrogated in other organisms including bacteria, *Drosophila melanogaster,* and human cell lines. While bacteria can upregulate DNA repair genes in response to DNA damage (*40*), enrichment of some of these transcripts in *Drosophila* and humans has been found to be at a typically modest level of only 1.5-3 fold (*20–25*). The level to which tardigrades enrich these transcripts, and the number of repair gene transcripts enriched, are by comparison far more extreme, and likely makes an important contribution to the extreme IR tolerance of some tardigrade species. We also found two histone subunits highly upregulated upon IR exposure (Table S1). Since we poly(A) selected our mRNA, these are likely to be poly(A)-containing mRNAs and hence non-cell cycle regulated histones that are typically used outside of S phase DNA replication cycles (*41*). We speculate that these strongly upregulated histones might contribute to forming new nucleosomes after DNA repair. Additionally, we identified that the DNA repair transcripts we investigated via *in situ* hybridization enrich in all or most tissues, and with more dramatic enrichment in certain tissues. The tissue specificity of enrichment did not suggest to us that different tissues use different upregulated DNA repair pathways (as an example, enrichment in ovary was seen for both *PCNA* (BER) and *XRCC5* (NHEJ)). It is currently unclear why these transcripts are enriched in a tissue-specific manner and if it is important to the ability of tardigrades to survive high levels of IR. A potential explanation for enrichment of these transcripts in secretory glands could be that secretory glands in general are more transcriptionally active than other tissues as they are specialized to make and export large amounts of protein (*34*). Transcriptionally active regions are more susceptible to DNA damage from IR (*42*), so potentially these tissues experience more DNA damage and upregulate the transcription of these genes disproportionately in response. However, we did not see evidence from our TUNEL experiments that these tissues experience more damage than others throughout the body.

Prior to this study, the only established mechanism of tardigrade IR tolerance was a protective mechanism that prevents damage, conferred by the *R. varieornatus* Dsup protein (*14–16*). Despite *H. exemplaris* having a Dsup protein (*15*), we found evidence for a response involving DNA repair genes rather than exclusively DNA protection. It is possible that the *H. exemplaris* Dsup does not have the same protective function as the *R. varieornatus* Dsup due to sequence divergence (*15*). We expressed both version of the *Dsup* gene in *E. coli*, and in this heterologous condition only the *R. varieornatus* Dsup protein conferred IR tolerance to bacteria (Fig. S6). The results of previous work (*14–16*, *43*) in combination with the results of this study suggest the possibility of synergy between protective and repair mechanisms in tardigrade IR tolerance. If some tardigrades use both mechanisms, the protective mechanism could work at IR levels at which DNA damage could be prevented or slow the accumulation of damage at higher IR levels, and as damage accumulates this could activate the transcription of DNA repair pathway genes that remedy the damage. Understanding how different mechanisms of IR tolerance might work together, as well as uncovering additional tolerance mechanisms, are intriguing avenues for future research.

## Supporting information

Protocol S1: Tardigrade TUNEL Protocol

Protocol S2: RNA isolation protocol

## Acknowledgments

We thank Dr. Nipam Patel and Dr. Jenny McCarthy for assistance with the adaptation of TUNEL protocols for tardigrades, the UNC High Throughput Sequencing Facility and the UNC Biology Microscopy Core for technical assistance, Dr. Dale Ramsden, Dr. Greg Matera, Dr. Tom Petes, Dr. Corbin Jones, Dr. Hemant Kelkar, Dr. Dan Janies, and members of the Ramsden and Goldstein labs for helpful feedback and discussions, and Adriana Schlachter for assistance with the CellProfiler analysis.

## Author contributions

CMCH and BG conceived of experiments. CMCH performed experiments and analyzed data. JDH conducted the analysis of bacterial expression level and protein solubility. TD performed trimming, mapping, and featureCounts analysis. CMCH, JDH, TD, and BG wrote and edited the manuscript.

## Data and materials availability

All data are available in the manuscript, the supplementary materials, or upon request.

## Materials and Methods

### Tardigrade culture

Cultures of *H. exemplaris* (Z151) were maintained as previously described (*45*). Animals were reared in 35 mm vented petri dishes (Tritech Research, T3500) with approximately 2 mL of Deer Park brand spring water and 0.5 mL Chloroccocum sp. algae (Carolina Biological Supply).

### Tardigrade irradiation

Gravid animals were collected and allowed to lay embryos and molt overnight. Freshly molted animals were placed into a 1.5 mL microcentrifuge tube in 100 µL of clean spring water (deer park) and then placed into a Gammator B Cs^137^ source gamma irradiator (current dose rate 1.4251 Gy/minute). Animals were left in the irradiator for an appropriate amount of time to reach the desired dose for each experiment (see below).

### Terminal deoxynucleotidyl transferase dUTP nick end labeling (TUNEL) assays, imaging, and analysis

Treated animals were irradiated as described above to a dose of either 2,180 Gy (24 hours) or 4,360 Gy (48 hours). Control animals were prepared in the same way as treated animals and remained on the lab bench for the same amount of time as their treated counterparts. Upon completion of irradiation, animals were either fixed immediately for TUNEL analysis or allowed to recover for 24 hours on the laboratory bench and then fixed. At least 20 animals were prepared for each treatment (not irradiated and irradiated 0 hr, and not irradiated and irradiated 24 hr). Fixation was performed in 4% paraformaldehyde (PFA) in PBS with 0.1% TritonX (0.1% PBT) overnight at 4°C. The fixative was washed out with 0.1% PBT. Animals were permeabilized by manual cutting with a syringe needle followed by sonication with a Branson Sonifier 250 probe sonicator (1 pulse, output control: 4, duty cycle: 50). The tardigrades were then transferred to a Mobicol column with 10 µm pore filter (Boca Scientific, M2210) in 0.1% PBT. Animals were subjected to a gradual methanol dehydration series from 25% to 100% methanol:0.1%PBT and left in −20°C to dehydrate overnight. Animals were gradually rehydrated from 100% to 0% methanol:0.1% PBT. The TUNEL assay protocol for tardigrades was adapted from a protocol for brine shrimp (McCarthy and Patel, personal communication) and for *Drosophila melanogaster* (*46*). Briefly, tardigrades were further permeabilized by incubating in Proteinase K (10 µg/mL) for 5 min at room temperature (RT) followed by incubations in *in situ* detergent (30 minutes, RT, shaking), 0.3% PBT Sodium deoxycholate (30 minutes, RT, shaking), and sodium citrate (1 hour, 65°C, shaking) (see Protocol S1 for details). TUNEL staining was performed as in (*46*) using the TMR red *in situ* cell death detection kit (Roche, 12156792910).

Stained animals were mounted in DAPI fluoromount-G (Southern Biotech) with 28.41 µm mounting beads (whitehouse scientific). Animals were imaged on a Zeiss 880 LSM with fast Airyscan detector. At least 9 individuals were imaged from each treatment for downstream analysis. 3D z-stacks were processed with FIJI (*47*) and saved as separated channel .tif files for processing in CellProfiler (*48*). Nuclei were segmented following the 3D segmentation of cell monolayer tutorial (tutorials.cellprofiler.org) through the “resize objects” as nuclei step. The mean intensity of TUNEL signal per nucleus was calculated in cell profiler using the “measure object intensity” module on identified nuclei in CellProfiler (*48*). Nuclear TUNEL intensity measurements were exported and the average “mean intensity” value was used for downstream analysis. Nuclear TUNEL intensity values were normalized to the mean of the not irradiated 0 hr samples for each experiment. One-way ANOVA followed by a post-hoc Dunnett test to the mean of the irradiated timepoint 0 for each experiment was used to determine significant differences between treatment groups.

### RNA sequencing

Approximately 200 adult animals were used for each replicate (3 replicates each of unirradiated, 100 Gy, 500 Gy, and 2,180 Gy). Animals were exposed to an appropriate dose of IR (or left on the lab bench for 24 hours, unirradiated) and RNA was isolated immediately from each replicate using the PicoPure RNA isolation kit (Applied Biosystems) following slightly modified manufacturer instructions (see Protocol S2). Libraries were constructed using the KAPA mRNA stranded library prep kit and fragmented to ~300bp. Paired end sequencing (2 x 50bp) was performed using the Illumina NextSeq2000 platform. Reads were adapter trimmed then mapped to the most recent genome for *H. exemplaris* (v3.1.5) using BBduk and BBmap (ver 39.01), and counts were assigned with featureCounts (ver 2.0.6) using the annotation file associated with this genome. Reads were aggregated at the level of genes and only genes with more than one count in at least two samples were kept for differential expression analysis. Transcript abundance, fold changes, and FDR values were determined using EdgeR (*49*).

### Gene homolog identification and cloning

Homologs of canonical DNA repair proteins of *H. exemplaris* were identified in a previous study (*50*) and updated to the current genome annotation (v3.1.5) using BLAST P. The homology of these proteins to their presumed DNA repair proteins was also confirmed by reciprocal BLAST to human and *Drosophila melanogaster* protein databases. A putative *H. exemplaris* Dsup protein was previously identified in (*15*). *H. exemplaris* POLQ was identified via BLAST P using human POLQ protein (NP_955452.3) as a query and confirmed via reciprocal BLAST. BARD1-like and BARD1-like C-terminal domain were identified via reciprocal BLAST. The partial Ku70 protein that is enriched upon exposure to IR (BV898_07145) was identified via an NCBI Domain search on the putative protein. This protein is predicted to only contain the N-terminal portion of Ku70 and lacks the domains responsible for interaction with Ku80 and DNA (*26*). Based on homolog transcript sequence, primers were designed to clone the full-length transcript from tardigrade cDNA or from GBlock synthesized gene fragments (IDT: *Rv Dsup, He Dsup, and RAD51*). Primers were designed with a 30bp overlap with the pDest17 expression vector (Invitrogen: 11803012) for subsequent incorporation into this vector via Gibson assembly.

### in situ hybridization and expression scoring

Templates for *in situ* hybridization probes were amplified from vectors containing the full-length gene using the primers listed in Table S5. Antisense RNA probes for *in situ* were synthesized as previously described (*51*, *52*), purified using an RNA clean and concentrator kit (Zymo, R1015), and eluted in RNAse free water. The final concentration of probes for *in situ* reactions was 0.5 µg/mL as previously recommended (*52*).

Tardigrades for *in situ* expression analysis were exposed to a dose of 100 Gy IR and fixed immediately for *in situ* hybridization as previously described (*53*). Controls were left on the lab bench for the equivalent amount of time. Fluorescent *in situ* hybridization was performed in adults as previously described (*53*). At least two replicates with 10 animals each were performed for each DNA repair transcript analyzed for both irradiated and control experiments. Samples were imaged on a Zeiss 880 LSM with fast Airyscan detector.

*in situ* hybridization expression profiles were examined in detail for at least 3 control and 3 treated individuals for each DNA repair transcript that we examined. Individuals were imaged at both lower laser power (appropriate setting for tissues with high expression) and higher laser power (to facilitate the observation of expression in tissues with lower levels of expression). Tissues were identified based on morphological analysis and informed by (*33*). Expression in each structure was scored from the higher laser power images on a scale from 0 (no observed expression) to 3 (observed oversaturated expression). A score of 1 indicates minimally observed expression and 2 indicates slightly undersaturated observed expression. Each tissue was scored within an individual and then a mean expression score for each tissue was calculated by averaging the tissue scores across individuals (Fig. S3).

### Bacterial protein expression and irradiation

pDest17 vectors containing full-length versions of individual tardigrade DNA repair transcripts were expressed in *E. coli* BL21 AI cells (Invitrogen, C607003) to determine if heterologous expression could confer tolerance to IR. The sequence of the expression vectors was confirmed before transformation into BL21 AI cells. Bacteria were grown overnight in 5 mL LB with Ampicillin and diluted 1:20 into LB with Ampicillin and 0.2% L-arabinose to induce expression from the pDest17 vector. Cultures were induced for 4 hours at 37°C while shaking. After 4 hours the OD600 of the cultures was measured. Induced cultures of bacteria expressing each DNA repair transcript were split into two 1.5 mL microcentrifuge tubes and densities were normalized by dilution into 1 mL total culture. Treated bacteria were exposed to a dose of 2,180 Gy IR while their control counterparts remained on the laboratory bench. After treatment, A dilution series of both treated and untreated bacteria was plated to determine the number of colony forming units (cfu). Percent survival was calculated as the cfu after irradiation divided by the cfu for untreated cells expressing the same DNA repair component. The percent survival was log transformed to standardize variance for statistical analysis as previously described (*54*). One-way ANOVA followed by a post-hoc Dunnett test to the means of the control groups (empty and GFP) was used to determine significant improvement in survival relative to controls following IR exposure.

### Analysis of heterologous protein expression in bacteria

Expression of protein in bacteria was analyzed by SDS-PAGE. Following the same induction protocol as used for irradiation experiments (see above), bacteria were pelleted by centrifugation, resuspended in 200 µL 0.85% NaCl, and lysed with a Branson Sonifier 250 probe sonicator (30 pulses, output control: 5, duty cycle: 50%). The soluble fraction of lysate was isolated by centrifuging at 14,000 rpm for 10 minutes at 4°C and retaining only the supernatant. Protein concentrations were quantified with Bio-Rad protein assay (Bio-Rad, 5000006).

Protein (2 µg total lysate, 5 µg soluble lysate) was loaded onto 4-12% Bis-Tris NuPAGE minigels (Invitrogen, NP0322BOX). 10 µL of precision plus protein kaleidoscope prestained standard (Bio-Rad, Cat#1610375) was included as a standard on each gel. Gels were run in 1x NuPAGE MOPS SDS running buffer (Invitrogen, NP0001) at 140 V for 75 minutes. Gels were stained in Coomassie and destained in a solution of 5:4:1 water:methanol:acetic acid before imaging (Fig. S4). The expected molecular weights of proteins that are reported were computed with the Expasy Compute pI/Mw tool (*55*).

### dsRNA synthesis and injection

DNA templates for synthesis of double-stranded RNA (dsRNA) for *XRCC5* and *GFP* were amplified using the primers indicated in Table S5. dsRNA for both *XRCC5* and *GFP* were synthesized as previously described (*36*), purified by isopropanol precipitation, and eluted in RNAse free water. dsRNA was diluted to a concentration of 1µg/uL in RNAse free water for injection. Gravid females for injection were isolated and allowed to lay eggs and molt overnight prior to injection. Adult tardigrades were injected with dsRNA targeting either *XRCC5* or *GFP* as previously described (*35*, *56*).

### RNAi Survival assays

Following injection with dsRNA targeting either *XRCC5* or *GFP* animals were allowed to recover overnight. After recovery, animals injected with either dsRNA were divided into two groups and placed into 1.5 mL microcentrifuge tubes. One group was exposed to 2,180 Gy IR and the other was left on the laboratory bench for an equivalent amount of time (24 hours). After treatment, animals were collected and placed into a 96-well plate with one animal/well filled with 100 µL spring water (Deer Park brand) and ~5 µL of chloroccocum algae (Carolina Biological Supply). IR exposure did cause some lethality on day 0 (the day animals were removed from the irradiator) in some groups (Fig. 4B). These animals were still transferred to 96-well plates along with surviving animals. Survival was checked approximately daily and the individuals in each well of the 96-well plate were scored as alive (movement detected) or dead (movement not detected) over the course of 7 days. This data was converted to survival curves in Prism and subjected to Kaplan-Meier simple survival analysis to determine significant differences in survival between groups (Fig. 4B).

Protocol S1. Detailed TUNEL protocol for *H. exemplaris*

Protocol S2. PicoPure RNA isolation protocol for *H. exemplaris*.

---

**Fig. S1.**
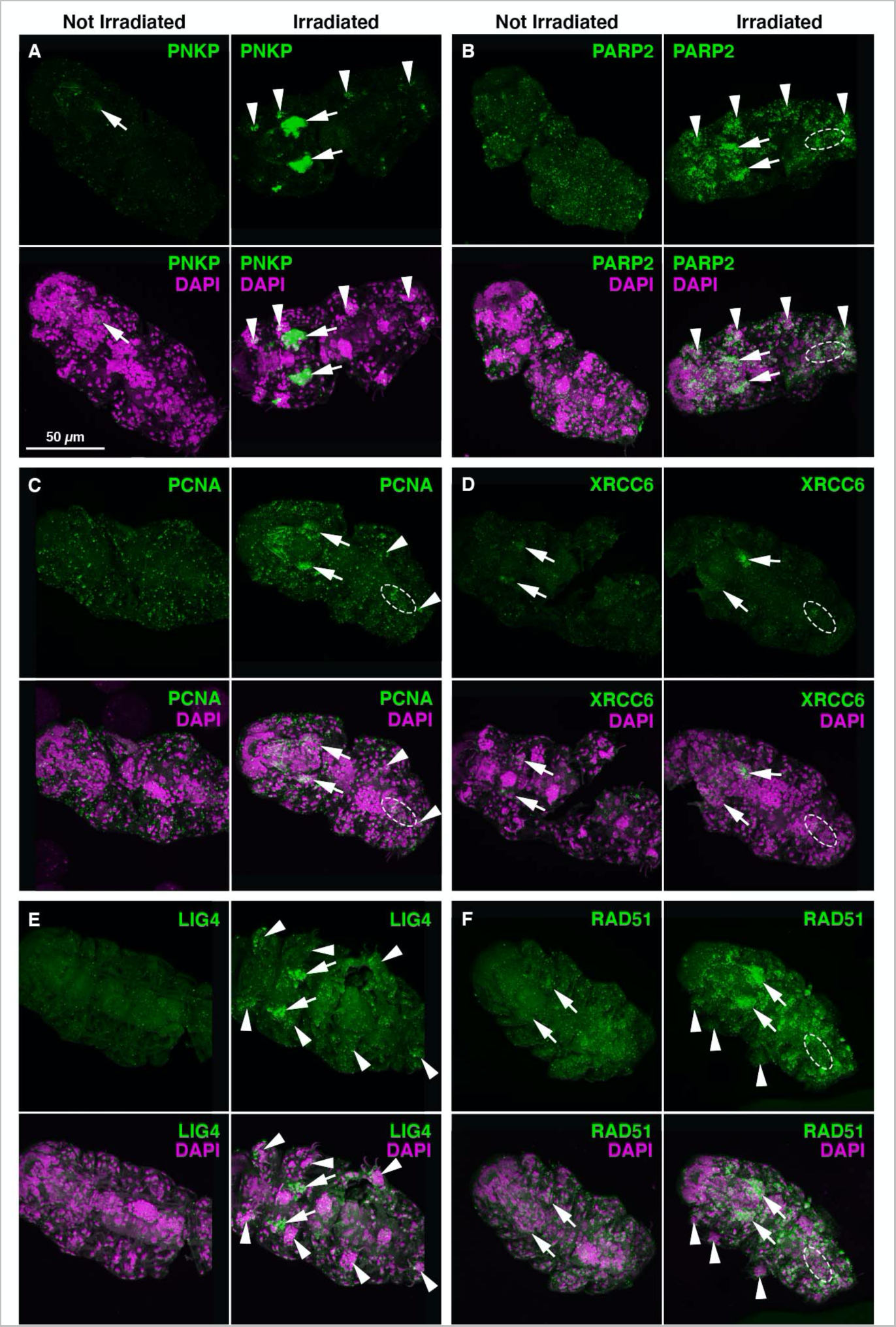
Tissue-specific enrichment of tardigrade DNA repair transcripts following ionizing radiation exposure. (**A-F**) Representative images of *in situ* hybridization for DNA repair transcripts with and without exposure to 100 Gy ionizing radiation. Exposure and contrast were adjusted here to visualize regions of most intense signal. Expression in salivary glands (arrows), claw glands (arrowheads), and hindgut (dashed outlines) is indicated where seen. Transcripts encoding members of the BER (**A-C**), NHEJ (**D-E**), and HR (**F**) pathways are represented. Scale bar in A applies to all images. Anterior is to the left.

**Fig. S2.**
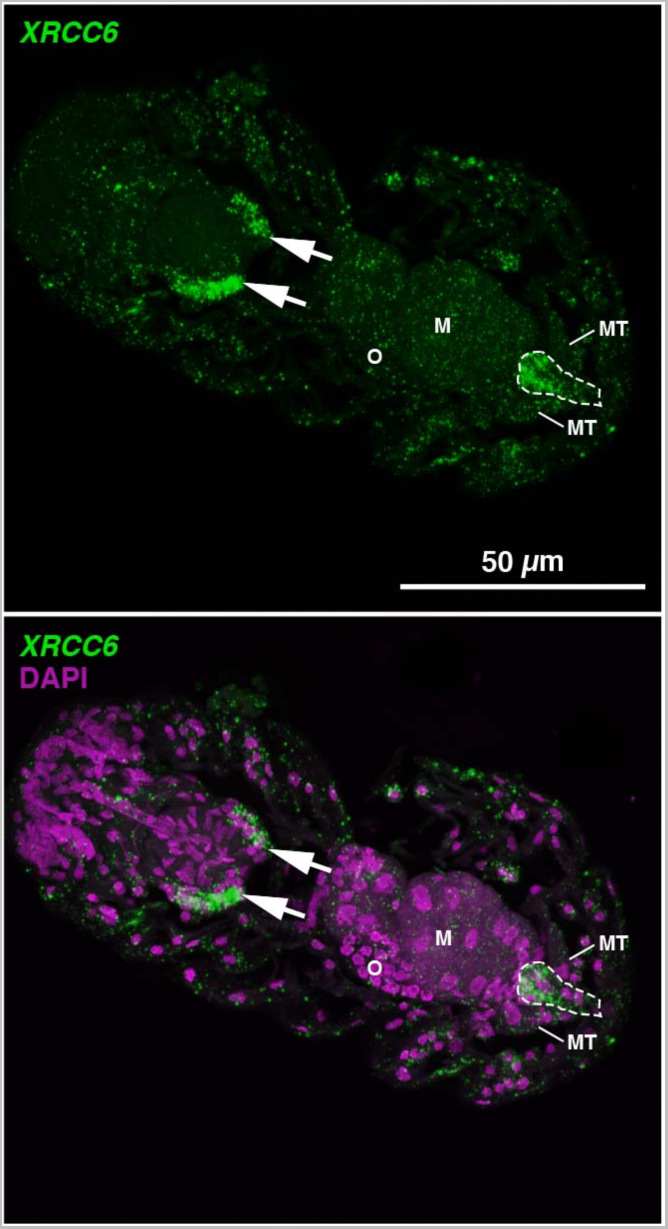
Enrichment of DNA repair transcript in hindgut of tardigrades following ionizing radiation. Maximum projection of optical sections containing hindgut expression of *XRCC6* to demonstrate hindgut location and identification. Expression in salivary glands (arrows) and hindgut (dashed outlines) is indicated where seen. Other landmark structures have been indicated as follows: O (Ovary), M (Midgut), and MT (Malpighian Tubules).

**Fig. S3.**
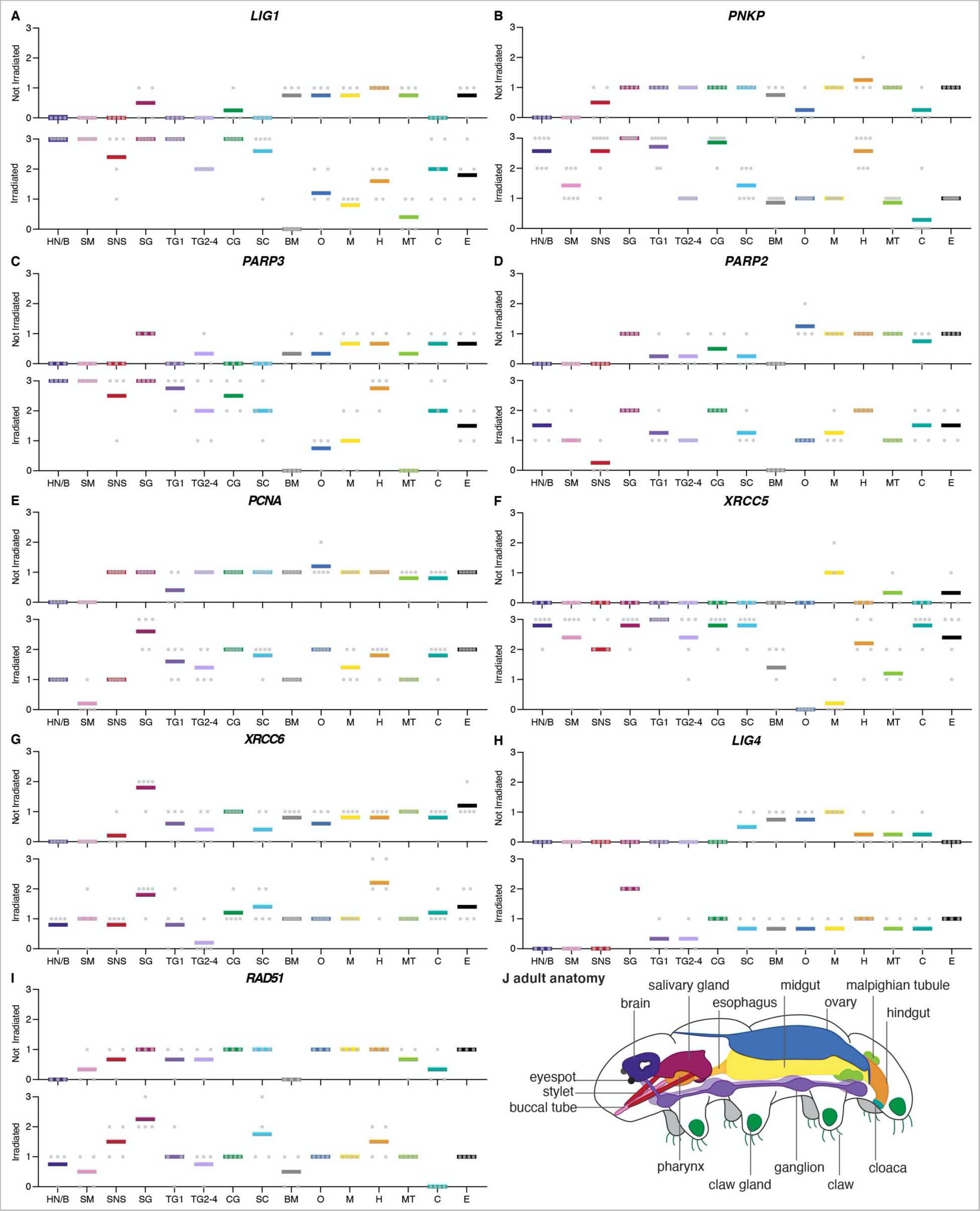
*in situ* hybridization for DNA repair transcripts reveals transcript accumulation in many tissues with some tissue-specific enrichment. (**A-I**) Tissue-specific enrichment profiles for DNA repair transcripts with and without exposure to 100 Gy IR. Tissue abbreviations are as follows: HN/B (Head Neuron/Brain), SM (Stylet Muscle), SNS (Stomodeal Nervous System, associated with stylet), SG (Salivary Gland), TG1 (Trunk Ganglion segment 1), TG2-4 (Trunk Ganglion segments 2-4), CG (Claw Gland), SC (Storage Cells, free-floating throughout the body cavity), BM (Body Muscle), O (Ovary), M (Midgut), H (Hindgut), MT (Malpighian Tubules), C (Cloaca), and E (Epidermis). Tissues were scored from 0 (no observed expression) to 3 (high expression) (see Materials and Methods for details on expression scoring). Tissue identification based on morphological analysis and informed by (*33*). Transcripts encoding members of the BER, TMEJ, NHEJ, and HR pathways are all represented. (**J**) Schematic of a lateral view of an adult tardigrade with landmark structures indicated (adapted from (*44*)).

**Fig. S4.**
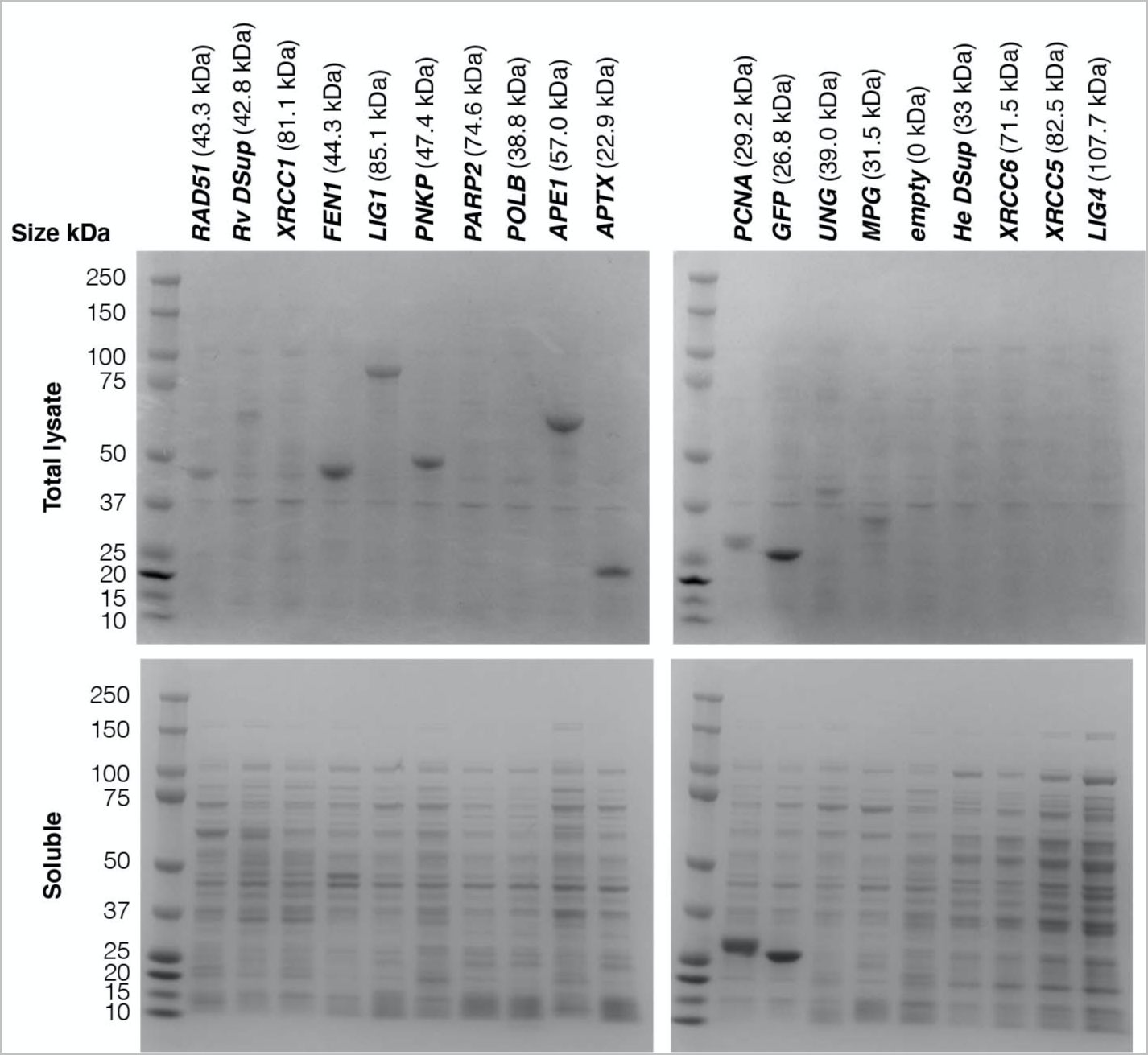
Bacterial expression of tardigrade DNA repair proteins. Protein gels showing levels of expression (top) and solubility (bottom) of tardigrade proteins heterologously expressed in *E. coli*.

**Fig. S5.**
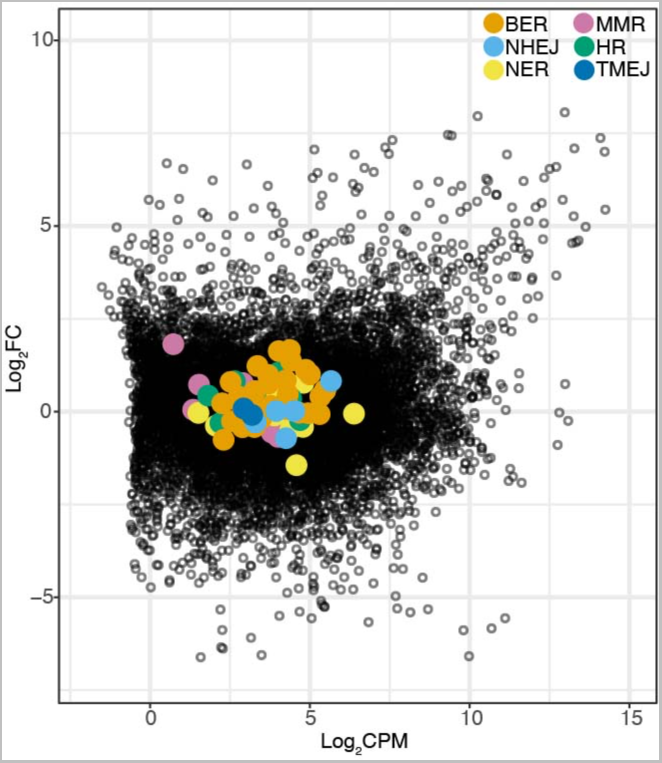
Transcriptional response of DNA repair genes to desiccation. Some BER and MMR transcripts are enriched slightly in response to desiccation. Original data from (*36*) and (*39*).

**Fig. S6.**
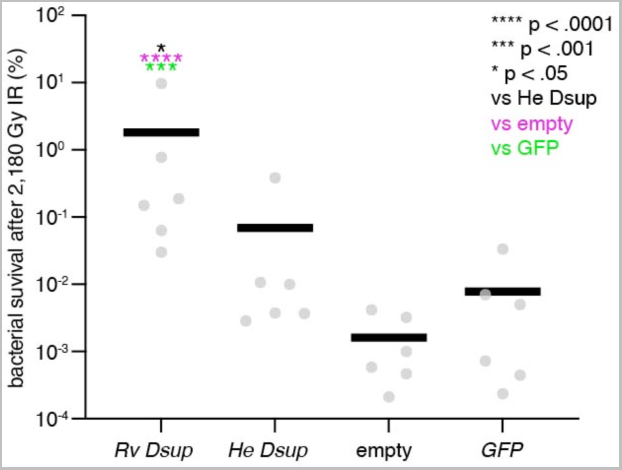
Expression of *R. varieornatus* Dsup but not *H. exemplaris* Dsup in bacteria improved bacterial survival after exposure to ionizing radiation. Controls are empty vector and a vector expressing GFP.

**Table S1.** Top 15 Significantly enriched transcripts following exposure to 500 Gy ionizing radiation. Transcripts that encode members of DNA repair pathways are highlighted in green.

| DNA Repair Pathway | Gene ID | logFC | logCPM | PValue | FDR | Protein |
| --- | --- | --- | --- | --- | --- | --- |
| N/A | BV898_10457 | 13.47446 | 9.7397723 | 0 | 0 | Hypothetical Protein: No conserved domains, tardigrade-specific |
| NHEJ | BV898_01166 | 8.3019188 | 12.811847 | 0 | 0 | XRCC5 (Ku80) |
| BER/TMEJ | BV898_18082 | 7.8223532 | 12.813553 | 0 | 0 | DNA Lig1 |
| N/A | BV898_03941 | 7.3494104 | 8.394104 | 0 | 0 | Hypothetical Protein - Histone H4 Domain |
| N/A | BV898_10478 | 7.1855765 | 10.834309 | 0 | 0 | Core histone macro-H2A.1 |
| BER | BV898_14774 | 7.0250053 | 9.9533286 | 0 | 0 | PNKP |
| BER | BV898_07590 | 6.7555331 | 11.95287 | 0 | 0 | PARP3 |
| N/A | BV898_17031 | 6.7823053 | 9.0346048 | 7.12E-320 | 1.27E-316 | Mitochondrial chaperone BCS1 |
| N/A | BV898_07145 | 6.3337674 | 7.9604761 | 3.40E-314 | 5.41E-311 | XRCC6 (Ku70) partial |
| N/A | BV898_09662 | 6.7606824 | 7.871877 | 2.01E-310 | 2.88E-307 | Hypothetical Protein: No conserved domains, tardigrade-specific |
| BER | BV898_08059 | 6.2385156 | 10.658468 | 3.26E-304 | 4.24E-301 | PARP2 |
| N/A | BV898_10564 | 9.5751369 | 6.3851308 | 2.05E-291 | 2.44E-288 | Protein phosphatase 1B |
| N/A | BV898_16497 | 6.7294928 | 8.2529798 | 1.86E-280 | 2.05E-277 | Hypothetical Protein - Ring and WWE domains |
| BER | BV898_09437 | 6.1884444 | 9.0754975 | 3.31E-265 | 3.38E-262 | PCNA |
| NHEJ | BV898_18536 | 5.3786796 | 9.7114364 | 5.27E-251 | 5.02E-248 | DNA Lig4 |

**Table S2.** Top 15 most highly represented transcripts after 500 Gy ionizing radiation. Ordered from highest-to-lowest mean log_2_TPM after 500 Gy IR exposure. Genes encoding housekeeping proteins and members of DNA repair pathways are highlighted in orange and green, respectively.

| Gene ID | Protein | 0 Gy Rep 1<br>log<sub>2</sub>TPM | 0 Gy Rep 2<br>log<sub>2</sub>TPM | 0 Gy Rep 3<br>log<sub>2</sub>TPM | 500 Gy Rep 1<br>log<sub>2</sub>TPM | 500 Gy Rep 2<br>log<sub>2</sub>TPM | 500 Gy Rep 3<br>log<sub>2</sub>TPM | 0 Gy Mean<br>log<sub>2</sub>TPM | 500 Gy Mean<br>log<sub>2</sub>TPM |
| --- | --- | --- | --- | --- | --- | --- | --- | --- | --- |
| BV898_08387 | Hypothetical Protein: No conserved domains | 15.29502578 | 15.22762843 | 15.16564047 | 15.31842213 | 15.27409912 | 15.15573694 | 15.22943156 | 15.24941939 |
| BV898_03848 | Elongation factor 1-alpha | 13.84955163 | 13.91029017 | 13.48685721 | 13.6739071 | 13.68387065 | 13.84198699 | 13.74889967 | 13.73325491 |
| BV898_04261 | 28S ribosomal RNA | 12.79454457 | 12.98725048 | 12.74634992 | 12.70987496 | 13.40741309 | 14.39251247 | 12.84271499 | 13.50326684 |
| BV898_16263 | SAHS 33020 | 13.5398077 | 13.69160557 | 13.804358 | 13.39478579 | 13.31782351 | 13.01263529 | 13.67859041 | 13.2417482 |
| BV898_02877 | Actin, cytoplasmic 1 | 14.05594588 | 14.17568206 | 13.70519287 | 13.0941487 | 13.10577203 | 13.33700564 | 13.97894027 | 13.17897546 |
| BV898_01166 | XRCC5 (Ku80) | 4.963104929 | 4.875845018 | 4.870232944 | 13.02542495 | 13.0097687 | 13.14600828 | 4.903060964 | 13.06040064 |
| BV898_07590 | PARP3 | 6.292615514 | 6.254191613 | 6.122859769 | 12.84059065 | 12.82790022 | 12.8760603 | 6.223222299 | 12.84818373 |
| BV898_17177 | putative Ovochymase-1 | 12.98240287 | 12.87385283 | 12.8654584 | 12.84017861 | 12.82080683 | 12.84905599 | 12.90723804 | 12.83668048 |
| BV898_01079 | Hypothetical Protein: No conserved domains, tardigrade-specific | 11.91406938 | 11.9486919 | 11.63996287 | 12.68759491 | 12.7734288 | 12.7219536 | 11.83424138 | 12.7276591 |
| BV898_18082 | DNA Lig1 | 5.028774136 | 5.035556912 | 4.912997025 | 12.6802133 | 12.63056783 | 12.73983375 | 4.992442691 | 12.6835383 |
| BV898_13380 | Hypothetical Protein: No conserved domains, tardigrade-specific | 12.77710543 | 12.76049904 | 12.2396206 | 12.50400969 | 12.51035015 | 12.64233128 | 12.59240836 | 12.55223037 |
| BV898_10202 | Hypothetical Protein: No conserved domains, tardigrade-specific | 12.39329356 | 12.30370182 | 12.15731514 | 12.51545016 | 12.5204354 | 12.59325264 | 12.28477017 | 12.54304607 |
| BV898_02951 | Hypothetical Protein: PTZ00491 super family | 8.186269453 | 8.35638015 | 8.339432709 | 12.49199281 | 12.43360992 | 12.25687412 | 8.294027437 | 12.39415895 |
| BV898_09382 | Cathepsin L1 | 12.31921247 | 12.27536324 | 12.38891681 | 12.35438508 | 12.29644179 | 12.29213972 | 12.32783084 | 12.3143222 |
| BV898_09692 | Hypothetical Protein: No conserved domains, tardigrade-specific | 7.989954268 | 8.132550376 | 7.781781809 | 12.28151443 | 12.27754942 | 11.92283828 | 7.968095484 | 12.16063404 |

**Table S3.** Additional significantly enriched DNA repair transcripts with log_2_FC > 3 following exposure to 500 Gy ionizing radiation.

| DNA Repair Pathway | Gene ID | logFC | logCPM | PValue | FDR | Protein |
| --- | --- | --- | --- | --- | --- | --- |
| HR | BV898_05956 | 5.4671337 | 10.057325 | 3.18E-232 | 2.39E-229 | BARD1-like |
| HR | BV898_00321 | 5.4364268 | 8.8928117 | 4.90E-225 | 3.34E-222 | Rad51 |
| HR | BV898_20143 | 5.3117773 | 7.7081581 | 1.76E-224 | 1.14E-221 | BARD1-like C-terminus domain |
| BER | BV898_11662 | 5.0021574 | 9.6958034 | 1.87E-222 | 1.17E-219 | XRCC1 |
| TMEJ | BV898_12022 | 5.7518622 | 10.02558 | 1.15E-219 | 6.51E-217 | DNA Pol Theta |
| NHEJ | BV898_13167 | 5.8565378 | 8.8210533 | 3.04E-209 | 1.61E-206 | XRCC6 (Ku70) |

**Table S4.**
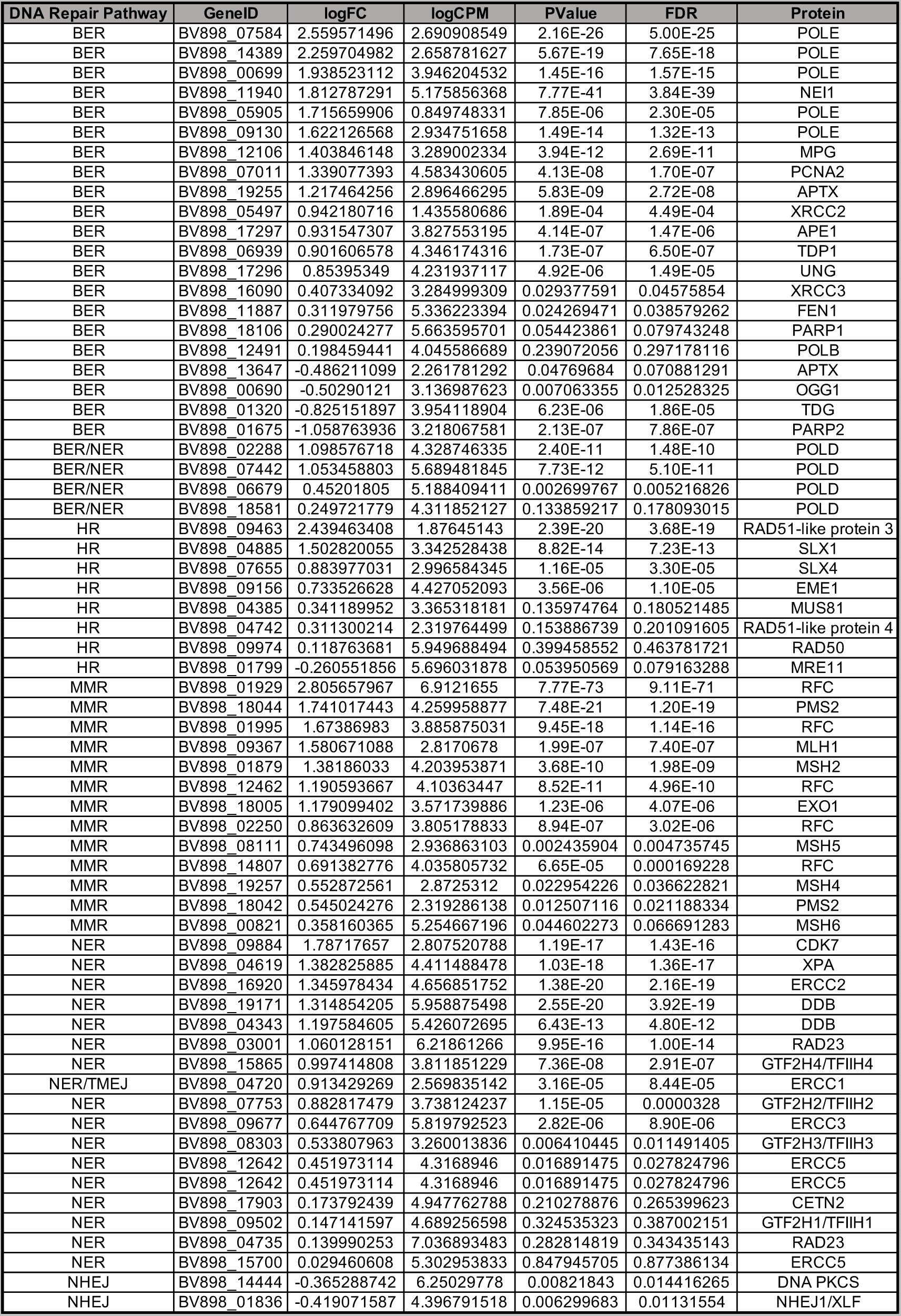
DNA repair transcripts that were not dramatically enriched following exposure to 500 Gy ionizing radiation.

**Table S5.**
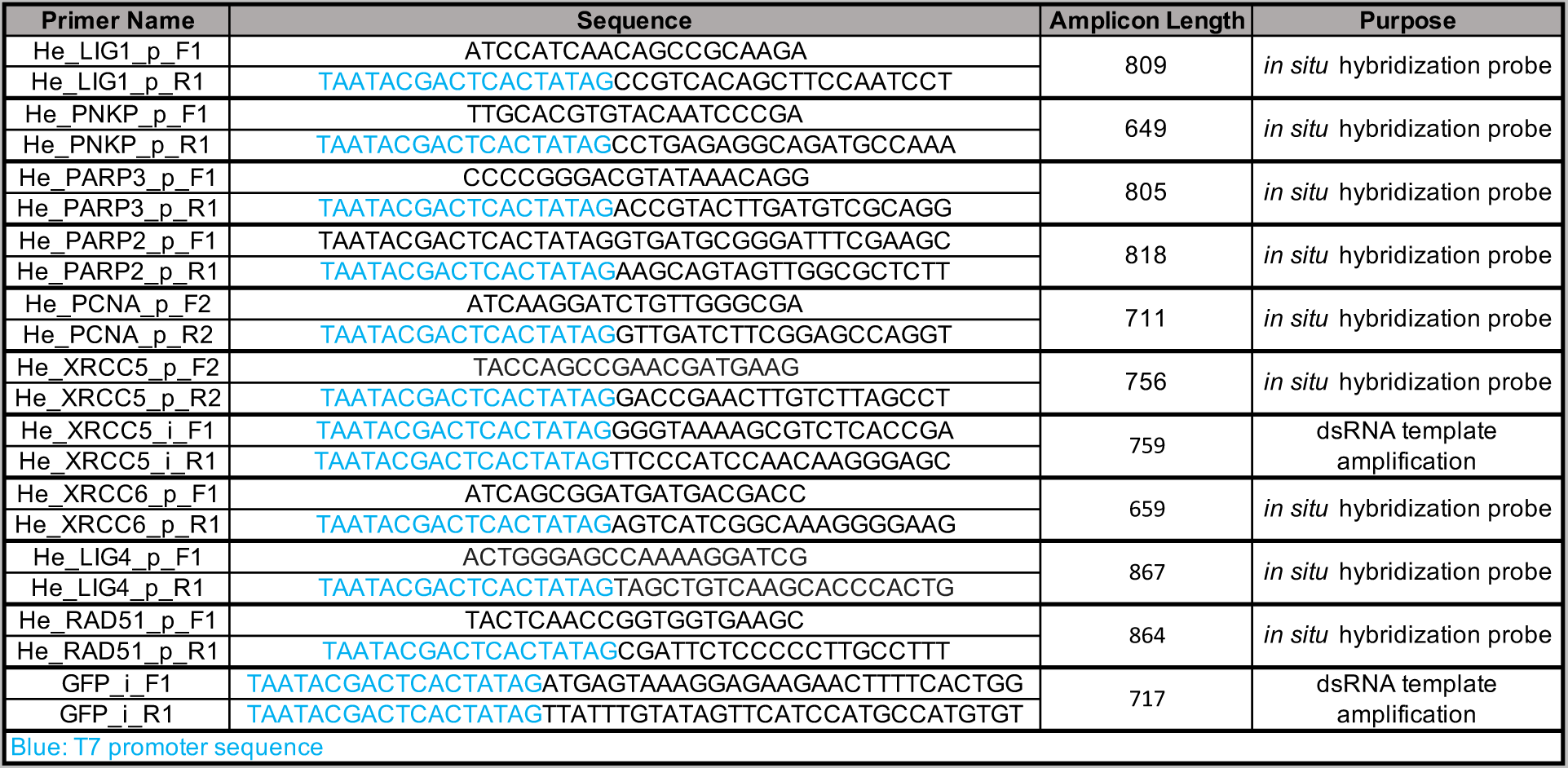
Primers used in this study.

