## Supplementary material for "Tardigrades dramatically upregulate DNA repair pathway genes in response to ionizing radiation": Protocol S1: Tardigrade TUNEL Protocol

***H. exemplaris* TUNEL protocol**

Fixation:

1. Spin animals down in 1.5mL microcentrifuge tube at 3000 rcf for 3 min
2. Remove water with syringe needle being careful not to disturb pelleted animals
3. Add 100µL seltzer water to the tube and wait 5 minutes to anesthetize animals
4. Make fix (4% paraformaldehyde (PFA) in 1x PBS with .1% Triton X (.1 % PBT)
5. Place animals in fixative and leave in 4°C overnight

Cutting and Sonication:

1. Wash out fix by spinning down animals at 3000 rcf for 3 min.
2. Remove fix with syringe needle being careful not to disturb pelleted animals
3. Replace with 100µL .1% PBT let sit for 5 min
4. Spin animals down at 3000 rcf for 3 min, remove .1% PBT
5. Repeat steps 3 and 4 for a total of 3 washes in .1% PBT
6. Using a glass Pasteur pipette transfer animals to a plastic petri dish with a small amount of .1% PBT
7. Using a syringe needle (26G) gently cut each animal in half
8. Return cut animals to 1.5mL microcentrifuge tube in .1% PBT
9. Spin cut animals down at 3000 rcf for 3 min
10. Sonicate animals using one pulse from a Branson Sonifier 250 probe sonicator set to output control 4, and duty cycle 50
11. Spin down sonicated animals at 3000 rcf for 3 min
12. Transfer sonicated animals to a petri dish with .1% PBT using a glass Pasteur pipette
13. Under a dissection scope, using a mouth pipette transfer the sonicated animal pieces into a prepared Mobiocol column with 10um pore filter (see below).

Column preparation:

1. Wash a prepared Mobiocol column with 10um pore filter (Boca Scientific, M2210) 5 x 5 min each with 500µL .1% PBT. Use a 1 mL syringe to push liquid through the column.

Methanol Dehydration:

1. Wash for 5 min each in 500µL 25%, 50%, 70%, and 90% methanol: .1% PBT
2. Wash 2 x 5 min in 500µL 100% methanol
3. Place in -20°C overnight

Methanol Rehydration

1. Wash for 5 min each in 500µL 90%, 70%, 50%, and 25% methanol: .1% PBT
2. Wash 3 x 5 min in 500µL .1% PBT

Permeabilization

1. Incubate in 500µL proteinase K solution (10µg/mL) for 5 min at room temperature (RT)
2. Wash 3 x 5 min in 500µL .1% PBT
3. Incubate in 500µL *in situ* detergent for 30 min RT shaking (750 rpm)
4. Wash 3 x 5 min in 500µL .1% PBT
5. Incubate in 500µL .3% PBT sodium deoxycholate for 30 min RT shaking
6. Wash 3 x 5 min in 500µL .1% PBT
7. Make sodium citrate solution fresh before use (1.47g in 50mL .1% PBT)
8. Incubate in 500µL sodium citrate for 1 hour at 65°C shaking
9. Wash 3 x 5 min in 500µL .1% PBT

TUNEL reaction (*in situ* cell death detection kit, TMR red, Roche, 12156792910)

1. Make TUNEL mix (90uL labeling mix, 10uL TdT enzyme)
2. Incubate samples in 100µL TUNEL mix for 3 hours at 37°C (shaking in the dark) (Use a syringe and associated stopper to keep the solution in the column)
3. Wash 5 x 5 min in 500µL .1% PBT at 37°C shaking
4. Mount with DAPI fluoromount-G (Southern Biotech) with 28.41 µm mounting beads (whitehouse scientific)

Solutions:

**.1% PBT:**

10x PBS	5 mL

10% TritonX	500 µL

ddH2O	to 50 mL

**Fixative:**

16% Paraformaldehyde	25 µL

.1% PBT	75 µL

**Proteinase K:**

1x PBS	2 mL

Proteinaske K (20mg/mL)	1 µL

**In situ detergent:**

10% SDS	5 mL

10% Tween 20	2.5 mL

1M Tris HCl (pH 7.5)	2.5 mL

.5M EDTA (pH 8.0)	100 µL

5M NaCl	1.5 mL

ddH2O	to 50 mL

**.3% PBT sodium deoxycholate:**

10% Triton X	1.5 mL

Sodium deoxycholate	.15 g

10% PBS	5 mL

ddH2O	to 50 mL
