## Supplementary material for "Tardigrades dramatically upregulate DNA repair pathway genes in response to ionizing radiation": Protocol S2: RNA isolation protocol

Tardigrade RNA isolation protocol (Using PicoPure Kit)

1. Turn on heat block and set to 42C
2. Collect tardigrades (100-200 animals) in RNAse Free tube (Fisher Scientific, 12-141-368)
3. Centrifuge at 4,500xG for 3 min to pellet the tardigrades
4. Remove all water and replace with 100uL Extraction Buffer (XB)
5. Using the blue pestle (that comes with tubes), disrupt the tardigrades – can check under microscope to make sure they are squished
6. Incubate at 42C for 30 minutes while shaking
   a. In the last 10 minutes of the incubation condition your column.
      i. Place 250uL Conditioning Buffer (CB) onto the column
      ii. Incubate at RT for 5 min
      iii. Centrifuge at 16,000xG for 1 min
   b. Toward the end of this incubation, pull out the DNAseI (Invitrogen) buffer from the freezer to thaw
7. After incubation, centrifuge your tube containing tardigrades at 3,000xG for 2 min
8. Pipette the supernatant containing the extracted RNA (100uL) into a new RNAse free tube, be careful not to disturb the pelleted tardigrade parts – do under microscope
9. Pipette 100uL 70% EtOH into your tube containing the RNA. Mix well by pipetting up and down multiple times
10. Pipette the mixture (200uL) into your preconditioned purification column
11. To bind RNA centrifuge for 2 min at 100xG immediately followed by a centrifugation at 16,000xG for 30s to remove flow through
12. Pipette 100uL Wash Buffer 1 (W1) into the purification column and centrifuge for 1 min at 8,000xG
13. Make DNAseI solution:
    a. 32uL RNAse Free H2O, 4uL 10x DNAseI Reaction Buffer, 4uL DNAseI
14. Pipette 40uL DNAse mixture onto the purification column and incubate at RT for 15 min
15. Pipette 40uL Wash Buffer 1 (W1) onto the purification column. Centrifuge at 8,000xG for 30s.
16. Pipette 100uL Wash Buffer 2 (W2) onto the purification column and centrifuge for 1 min at 8,000xG
17. Pipette another 100uL Wash Buffer 2 (W2) onto the purification column and centrifuge at 16,000xG for 2 min. Discard flow through.
18. Centrifuge 16,000G x 1 min to dry column
19. Transfer column to new .5mL centrifuge tube provided in the kit
20. Pipette 11uL of Elution Buffer (EB) directly onto the membrane of the purification column
21. Incubate the column for 1 min at RT
22. Centrifuge the column for 1 min at 1,000xG to distribute the EB and then spin for 1 min at 16,000xG to elute the RNA
23. Check concentration of RNA on nanodrop
24. Use immediately or store at -80C
